# Easy identification of insertion sequence mobilization events in related bacterial strains with ISCompare

**DOI:** 10.1101/2020.10.16.342287

**Authors:** E.G. Mogro, N. Ambrosis, M.J. Lozano

## Abstract

**Motivation:** Bacterial genomes are composed by a core and an accessory genome. The first composed of housekeeping and essential genes, while the second is composed, in its majority, of mobile genetic elements, including transposable elements (TEs). Insertion sequences (ISs), the smallest TEs, have an important role in genome evolution, and contribute to bacterial genome plasticity and adaptability. ISs can spread in a genome, presenting different locations in nearly related strains, and producing phenotypic variations. Few tools are available which can identify differentially located ISs (DLIS) on assembled genomes.

**Results:** We developed ISCompare to profile IS mobilization events in related bacterial strains using complete or draft genome assemblies. ISCompare was validated using artificial genomes with simulated random IS insertions and real sequences, achieving the same or better results than other available tools, with the advantage that ISCompare can analyse multiple ISs at the same time and outputs a list of candidate DLIS. We think that ISCompare provides an easy and straightforward approach to look for differentially located ISs on bacterial genomes.

**Availability and implementation:** ISCompare was implemented in python3 and its source code is freely available for download at https://github.com/maurijlozano/ISCompare.

**Supplementary information:** Supplementary data are available at https://github.com/maurijlozano/ISCompare.

## 1 Introduction

Bacterial genomes are a mosaic composed of a core genome of housekeeping and essential genes, and an accessory genome, composed, in its majority, of mobile genetic elements, which can be grouped into two classes, plasmids and bacteriophages, and transposable elements (TEs) (Siguier *et al*., 2014). Insertion sequences (ISs) are the smallest TEs, being only formed by imperfect terminal inverted repeats and a transposase coding region (Siguier *et al*., 2014; Mahillon and Chandler, 1998). Once an IS is acquired it can spread in a genome by transposition, and though most insertions might be deleterious, some may be beneficial and confer an increased fitness or provide a selection advantage (Schneider and Lenski, 2004). One of the most important roles of ISs is their participation in genome evolution, being mediators of genomic rearrangements, gene inactivation, over-expression and modulation of the expression of neighbor genes (Siguier *et al*., 2014). ISs contribute to bacterial genome plasticity and to the adaptability of its phenotypic traits (Schneider and Lenski, 2004), such as resistance to antibacterial agents, virulence, pathogenicity, catabolism (Vandecraen *et al*., 2017) and defence against harmful genes (Fan *et al*., 2019). IS insertions have been demonstrated to participate in the adaptation of bacterial strains to antibiotic treatments (Mugnier *et al*., 2009) and vaccination strategies (Pawloski *et al*., 2014; Carriquiriborde *et al*., 2019; Zomer *et al*., 2018). For these reasons, profiling IS insertion sites and its variation between related strains has gained importance.

The principal methods used for IS profiling have been chromosomal DNA hybridization (Fan *et al*., 2019; Soria *et al*., 1994), restriction fragment length polymorphism (Das *et al*., 1995) and a diversity of PCR related methods (Bik *et al*., 1996; Lozano *et al*., 2010; Suzuki *et al*., 2004). With the advent of high throughput sequencing technologies, whole genome sequences have become readily available for most microorganisms, and the ISs locations can be found using several tools such as ISFinder (Siguier *et al*., 2006), ISEScan (Xie and Tang, 2017), Oasis (Robinson *et al*., 2012) and ISQuest (Biswas *et al*., 2015). Further, some tools for the comparison of IS location between bacterial strains have been developed, most of them based on the soft mapping of short reads from whole genome shotgun sequencing experiments to a reference genome (Breseq, Barrick et al., 2014; Transposon Insertion Finder, Nakagome et al., 2014; ISMapper, Hawkey et al., 2015; and panISa, Treepong et al., 2018). Some disadvantages of these programs are that they require a high genome coverage for an accurate detection of ISs, and that they are difficult to apply to the analysis of massive genomic datasets (Adams *et al*., 2016). Also, laboratories in developing countries often do not have the resources to sequence many bacterial isolates with the required sequencing depth, which makes the use of the previous programs difficult.

A complementary tool, ISseeker (Adams *et al*., 2016), was designed for the rapid and high-throughput mapping of ISs using whole genome sequence assemblies. A special case are draft genome assemblies, where contigs are typically broken at IS locations (Sohn and Nam, 2018), containing partial IS sequences in one or both ends. This tool identifies the locations of a provided insertion sequence using blast (Altschul *et al*., 1990), then its flanks are extracted and mapped against a reference genome. However, the comparison to establish differentially located ISs (DLIS) has to be manually done. Additionally, the analysis using ISseeker requires some background knowledge of the ISs present on the test organisms, and it can only search for one IS at a time.

In order to automate the detection of IS mobilization events in related bacterial strains we developed a new program, ISCompare, which uses complete or daft genomes as query and reference, to find DLISs. The method is freely available in the form of opensource code, and here we validate its use via the analysis of simulated and real genomic sequences.

## 2 Algorithm and Implementation

### 2.1 ISCompare

An overview of ISCompare workflow is shown in figure 1. ISCompare was written in python and uses blastn (Altschul *et al*., 1990), and Biopython (Entrez and SeqIO, Cock et al., 2009), DNA_features_viewer (Zulkower and Rosser, 2020), Pandas (Reback *et al*., 2020) and Numpy (Van Der Walt *et al*., 2011) python modules. ISCompare takes as input query and reference genomes in genbank flat format or its corresponding accession numbers, both of which can be either complete or draft genomes. If a list of accession number is provided, ISCompare will download the genomes sequences from NCBI Assemblies (Kitts *et al*., 2016). An optional multifasta DNA file containing the ISs database can be supplied, otherwise IScompare can search for the ISs present through the use of ISFinderBlast.py script (-I option). This script uses the mechanize (Mechanize – Automate interaction with HTTP web servers. v0.4.5) python module to launch an IS search at ISFinder webpage (Siguier et al., 2006; http://www-is.biotoul.fr). Then it collects the sequences of the found ISs into the IS.fna file located on the results folder. In the first step a blastn search of the query genome is run against the reference genome. This step aims at removing complete identical scaffolds or replicons from the subsequent analysis. A sequence will be considered identical if it has a qcovhsp (i.e. query cover per high-scoring pair) greater than 99% and less than 20 unaligned nucleotides (Fig. 1. Step 1). With the remaining query sequences the program searches for ISs by running a blastn search against the IS database. Here, blastn is run with culling_limit option set to 1 to avoid redundant hits (i.e. hits from the same genomic region with different ISs on the database), and a settable E-value cutoff (1.10^−10^ by default). IS hits with an alignment length smaller than a third of the detected IS length, except for hits on scaffold ends (which on draft assemblies usually contain partial ISs), are removed. Additionally, small query scaffolds corresponding mostly to an IS hit are removed from the analysis. Next, the sequences adjacent (500 base pairs on each side by default) to the detected ISs are extracted (<u>Q</u>uery <u>I</u>S <u>F</u>lanks - QIF), concatenated, exported as a multifasta file, and blasted against the IS database. If an IS is found on a QIF, its name is compared to the original IS, and if it is related (i.e. same IS, same group or same family), the QIF is tagged for manual verification since adjacent related ISs may produce false positives (Supplementary material, Fig. S5. Panels E, G, H, J).

**Fig. 1.**
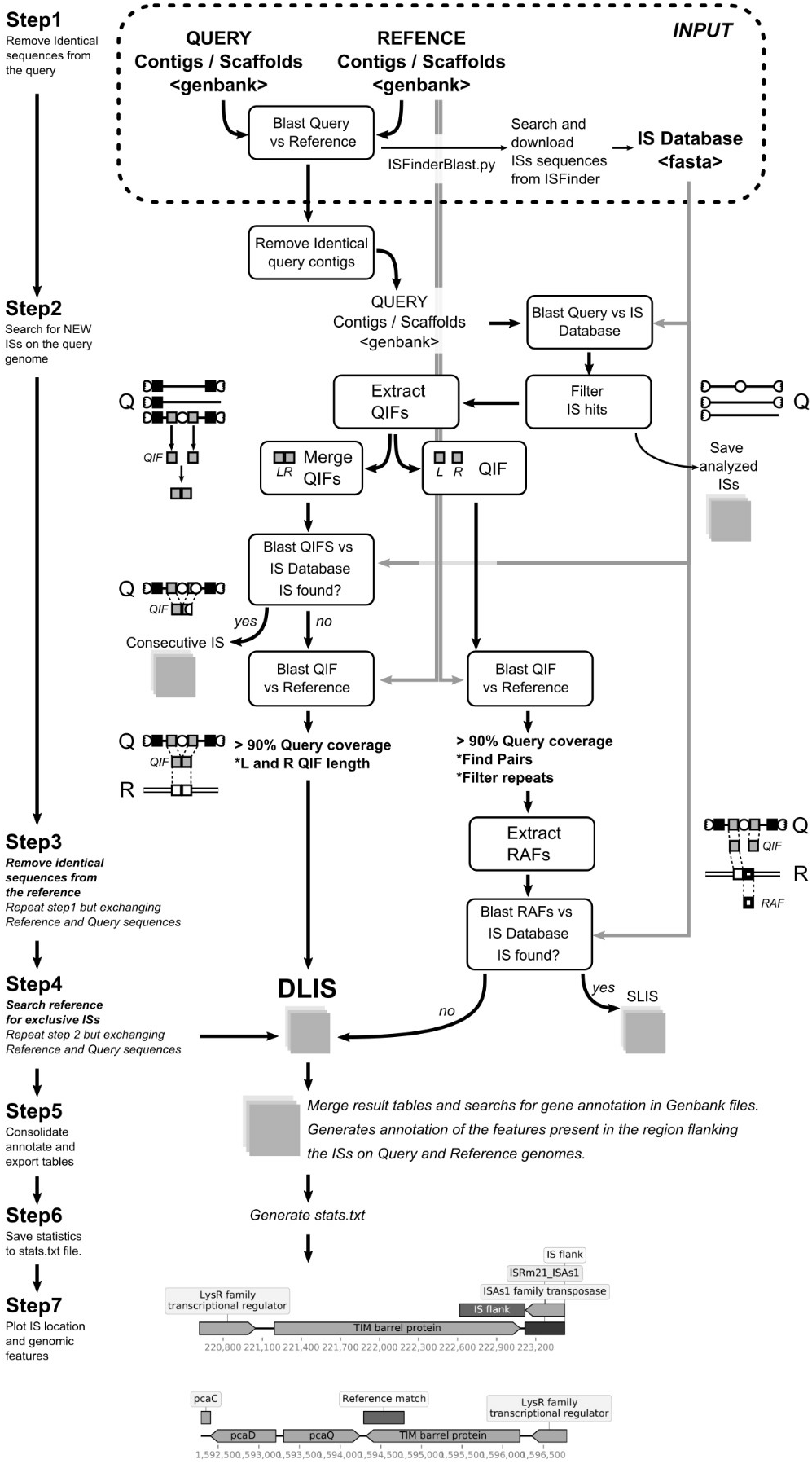
Outline of ISCompare algorithm. Schematic representation of the steps involved in the determination of differentially located insertion sequences by ISCompare. Rounded boxes represent processes. White circles: ISs. Black and grey boxes: query IS flanks (QIFs). Thick edge boxes: reference anchor sequence flanks (RAFs). Q: query. R: reference.

The following step makes a blastn search of the concatenated QIFs (only those containing at least a fifth of the flank length from each side of the IS) against the reference genome, and looks for hits with high query coverage (>90%). The hits meeting this criterion correspond to ISs only present in the query genome (Fig. 1, Step 2).

Next, QIFs are extracted independently and blasted against the reference genome. The hits with a query coverage greater than 90% are used as anchors to look for the presence or absence of ISs within the neighboring genomic regions in the reference genome (<u>R</u>eference <u>A</u>nchor <u>F</u>lanks - RAF). At this step only the RAF where an IS is expected (upstream or downstream) are analysed (Fig. 1, Step 2). Here several cases may arise. Some of the QIFs will match to two segments of the reference genome separated either by the length of an IS (i.e. both the query and reference genome have the IS in the same location) or by a smaller distance (i.e. the IS is only found on the query genome). In addition, if some QIFs contain repeated sequences, more than two matches with the reference genome will be produced. In those cases, the software tries to find the correct pairs by analysing their location in the reference genome. If such a pair is formed, the QIFs are analysed, else they are tagged as unspecific. In the case of draft assemblies, most ISs are located on scaffold ends, thus QIFs will have sequences from one IS flank only. In those cases, blastn searches against the reference genome will produce (ideally) only one hit. If more than one hit is produced, the QIF will be tagged as unspecific. Next, the corresponding RAFs are extracted from the reference genbank file and blasted against the IS database. If an IS of the same class or family is found, it means that the same insertion is present on both genomes, otherwise, an IS exclusively found in the query genome is informed.

In the following step, the same procedure is repeated but using the query genome as reference and the reference genome as query. This analysis will report ISs only found in the reference genome. All the results are consolidated in a single table containing the reported DLISs, the location of QIFs and reference anchor regions, and the annotation information from both the query and reference genomes. Finally, DNA_features_viewer module (Zulkower and Rosser, 2020) is used to plot a pdf file with schematics of the reported ISs location and surrounding genes. For some applications, it is useful to move the QIFs position away from the detected IS. The shift option (-S) can be used in such cases to increase the size of the region detected as an IS in a specified number of nucleotides, shifting the QIFs away (Supplementary material, Fig. S6.A).

The program can be run on computers with low resources. All the tests were run on common laptop computers, and on a virtual PC running with three CPUs and 3GB of RAM. For command line options see the ISCompare github page (https://github.com/maurijlozano/ISCompare).

### 2.2 ISsimulator

In order to generate simulated genome sequences with random IS insertions, a new script ISsimulator.py, was used. ISsimulator was written in python and requires Biopython (Cock *et al*., 2009), a reference genome in genbank format and the IS to be inserted in fasta format. The program inserts a selected number of copies of the insertion sequence at randomly chosen genomic locations, and outputs modified genbank and fasta files, and a table with the location of the inserted ISs.

### 2.3 Sequences used on this work

All the genome sequences used on this work were downloaded from the NCBI or EMBL-EBI genome databases. The corresponding accession numbers are listed in supplementary table S1. A compilation of IS sequences downloaded from https://github.com/thanhleviet/Isfinder-sequences was used as IS database, except in the cases where ISCompare was used to look for a single IS.

### 2.4 Average Nucleotide Identity and digital DNA-DNA hybridization analysis

Average Nucleotide Identity (ANI) and digital DNA-DNA hybridization (dDDH) analysis were performed at the ANI/AAI-Matrix Genome-based distance matrix calculator (Goris et al., 2007, http://enve-omics.ce.gatech.edu/g-matrix/) and the GGDC Genome-to-Genome Distance Calculator 2.1 (Auch et al., 2010, http://ggdc.dsmz.de/ggdc.php#) web servers. Phylogenetic trees were visualized with ITOL (Letunic and Bork, 2019) and when necessary edited with Inkscape (v1.0, inkscape.org).

### 2.5 Statistical analysis and figures

All the statistical analysis and the corresponding plots were done in R software (Team, 2014). Final figures were edited with Inkscape (v1.0, inkscape.org).

## 3 Results

### 3.1 ISCompare parameter optimization

In order to optimize the program parameters we generated artificial *Eschericia coli* str. K-12 substr. MG1655 and *Ensifer meliloti* 2011 genomes with 100 random IS30 or ISRm5 insertions correspondingly, and run ISCompare using different settings in a stepwise mode (Supplementary material, Table S2). We made two kinds of comparisons: in the first case, we compared the *E. coli* str. K-12 substr. MG1655 genome with the artificially generated one (i.e. identical genetic background); in the second case we compared *E. meliloti* 1021 with the artificially generated *E. meliloti* 2011-100IS genome. *E. meliloti* 1021 and *E. meliloti* 2011 are very closely related but not identical strains, which make them good as testing subjects (Supplementary material, Fig. S1).

ISCompare was run with different surroundingLen [-s option] (i.e. the length of sequence to extract from each side of a detected IS) values ranging from 100 to 2000 nucleotides in steps of 100. ISCompare outputs a results table with the detected differentially located ISs under the ‘DLIS’ category for the most confident cases, and within several categories of ‘Verify manually...’ or ‘Discarded from the analysis...’ cases (Supplementary material, Fig. S5; VD for now on) which may result in a false positives, as a consequence of repeated genomic regions, consecutive identical (or very similar) ISs or when there is not a significant blastn hit for the QIFs on the reference genome. In the comparison of *E. coli* K12 genomes, where the genetic background is identical, a precision of 100% (93%) and a sensitivity of 91% (99%) were achieved for surroundingLen in the range of 200-500 nucleotides for the DLIS category. Values in parenthesis were obtained by taking the VD categories as DLIS (Supplementary material, Fig. S2). When the *E. meliloti* strains were compared, it was not possible to achieve a precision and a sensitivity greater than 90% for both DLIS and VD categories for a single surroundingLen. For DLIS results, the best surroundingLen was in the range of 400-1100 nucleotides with a precision of 98.9% and a sensitivity 94%, while for VD the best surroundingLen was in the range from 500 to 2000 nucleotides (Supplementary material, Fig. S2) with the best results for 1400 with a sensitivity of 100% and a precision of 86%.

For the optimization of the remaining parameters, a surroundingLen of 500 nucleotides was used as it produced, in general, good results in all the analysed cases.

Using the optimal surroundingLen the minAlnLength (i.e. the minimal required alignment length of the QIFs to the reference genome), minlength (i.e. the minimal QIFs length to be extracted from the genome genbank file), ISdiff (i.e. the minimal difference between the blast alignment length and the query length, which is used to discard scaffolds that only match to IS sequences) and scaffoldDiff (i.e. the maximum number of allowed nucleotides missing on the blastn alignment between the query and reference genomes) were optimized by manually assigning different values. The minAlnLength and minlength arguments are more relevant for the analysis of draft genomes, where in the case of small contigs, there might be QIFs with high query coverage, but with a length smaller than surroundingLen. Minlength option is used in such cases to filter QIFs with length minor to minlength, while minAlnLength is used to filter QIFs with a blastn alignment to the reference genome of a length smaller than minAlnLength. To optimize these parameters we searched for draft genome assemblies nearly related to *E. meliloti* 1021, and compared them with the reference *E. meliloti* 1021 strain genome (Supplementary material, Fig. S3). *E. meliloti* strain USDA1022 was used (Supplementary material, Table S1). For the assessment of true positives, the results were manually revised since no previous reports were made on the IS distribution on this strain (Suplementary table S2). The optimal values (minAlnLength=50, minAlnLength=50, ISdiff=50 and scaffoldDiff=20) were established as the default for the program.

### 3.2 Detection of IS location changes using artificially generated genomes

To assess the sensitivity and precision of our software we compared the *E. coli* str. K-12 substr. MG1655 genome with artificial genomes on which random IS insertions were made using ISsimulator script. In addition, we used an IS database containing all the insertion sequences found by ISFinder web server. A total of 3000 IS insertions of IS30 were simulated in steps of one hundred per genome and analysed using ISCompare (i.e. the program was run 30 times). Our method demonstrated to have a very good recall (94%) and precision (100%), with no false positives and few false negatives (181 over 3000 positives) when analysing the DLIS category (Supplementary material, Table S3). The program almost reached 100% recall (99.1%) at the expense of precision (92%) when considering as true the VD categories. It should be noted that in the case of *E. coli* str. K-12 substr. MG1655, on average every 100 differential ISs 12 sequences were tagged for manual inspection, which could be easily done with the -p option producing a graphic pdf report of the IS genomic context on both query and reference genomes. In general, the ISs tagged for manual inspection represented false positives cases (40%) mostly due to consecutive ISs (Supplementary material, Table S3).

IScompare can be also run to look for a specific IS (Supplementary material, Table S3). In such a case the results for the simulated data were similar. This was expected due to the capacity of ISCompare to distinguish consecutive insertions of different ISs.

### 3.3 Comparison of *E. meliloti* 1021 and *Pseudomonas aeruginosa* PAO1 with strains with progressively decreasing digital DNA-DNA hybridization

In order to test ISCompare with real genomic data, we selected different strains of *P. aeruginosa* with complete closed genomes and with progressively decreasing values of digital DNA-DNA hybridization (dDDH). Ten strains meeting this criterion were selected and *P. aeruginosa* PAO1 was used as reference (Supplementary material, Fig. S4 and Table S4). The results were manually analysed to assess for true/false positives cases. Only the cases where an IS was found either on the reference or query genomes, and which flanks were perfectly conserved (i.e. equal genomic context) were considered as true positives. On average, 4.6 DLIS were found on each genome (6.3 for the VD reports), with a maximum of 9 and a minimum of 0. The precision was very good (100%) for the DLIS category, but was greatly diminished (29%) when the VD cases were taken as positive (Supplementary material, Table S4). It should be noted that most of the VD cases are only reported with the objective of improving the sensitivity after a manual inspection, since some true positives are usually tagged for manual verification due to repetitions and consecutive ISs. A significant correlation between dDDH and the number of discarded QIFs was found (*R=−0.82, p=0.012*), since, as it was expected, more QIFs were discarded in the filtering steps when the compared genomes were phylogenetically more distant. The distribution of the DLIS mapped to the reference genome is shown in figure 2.A.

**Fig. 2.**
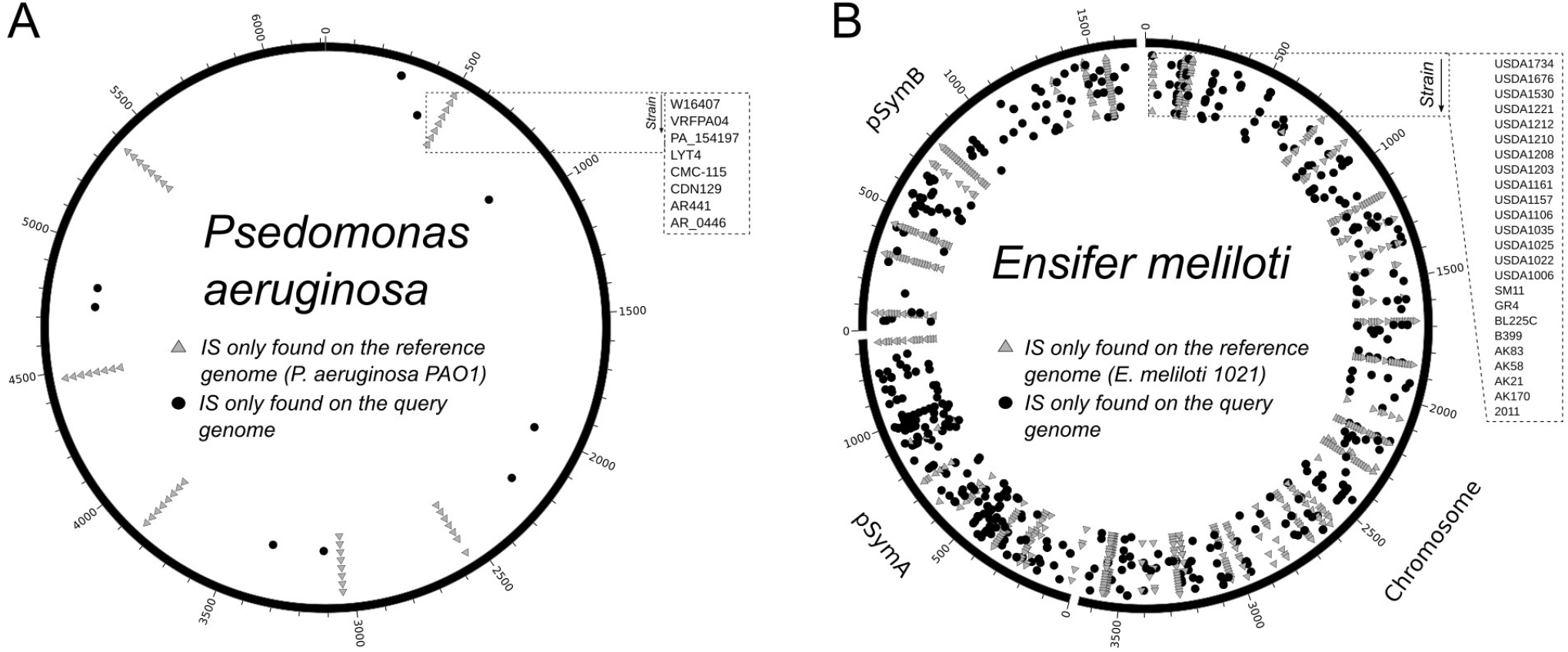
Genomic distribution of DLIS. Circos plot showing the genomic location of DLIS in *P. Aeruginosa* and *E. Meliloti* strains. A. DLIS found in *P. aeruginosa. P. aeruginosa* PAO1 was used as reference and compared to strains AR_0446, AR441, CDN129, CMC-115, LYT4, PA_154197, VRFPA04 and W16407, shown in that order from the border to the center. B. DLIS found on *E. Meliloti. E. Meliloti* 1021 was used as reference and compared to strains 2011, AK170, AK21, AK58, AK83, B399, BL225C, GR4, SM11, USDA1006, USDA1022, USDA1025, USDA1035, USDA1106, USDA1157, USDA1161, USDA1203, USDA1208, USDA1210, USDA1212, USDA1221, USDA1530, USDA1676 and USDA1734, shown in that order from the border to the center. Light-gray triangles: DLIS found on the reference genome. Black circles: DLIS found on the query strains.

As very few DLIS were found in the case of *P. aeruginosa*, ISCompare was used to identify DLIS in *E. meliloti*, which is known to have a greater number of ISs (Sallet *et al*., 2013). Twenty six *E. meliloti* strains were compared to *E. meliloti* 1021 using ISCompare (Supplementary material, Fig. S1, Fig. S3, Table S1 and Table S5). The results were manually analysed to assess for true/false positives cases, since no previous report comparing the distribution of insertion sequence in these strains could be found. We considered as true positives only the cases where the genomic context was the same on both sides of the IS. Our software found an average of 40 DLIS on *E. meliloti*, with values fluctuating between 0 (strain 1021-2119 comparison) and 86 (strain 1021-USDA1212) (Fig. 2.B). Most of the DLIS were located on the symbiotic plasmid pSymA (0.194 DLIS/Kb), followed by the chromosome (0.172 DLIS/Kb) and the chromid pSymB (0.134 DLIS/Kb), although no significant difference was observed (Fig. 3). That the pSymA presented a higher frequency of mobilization events was expected, in accordance with the higher plasticity associated with plasmids (López *et al*., 2019).

**Fig. 3.**
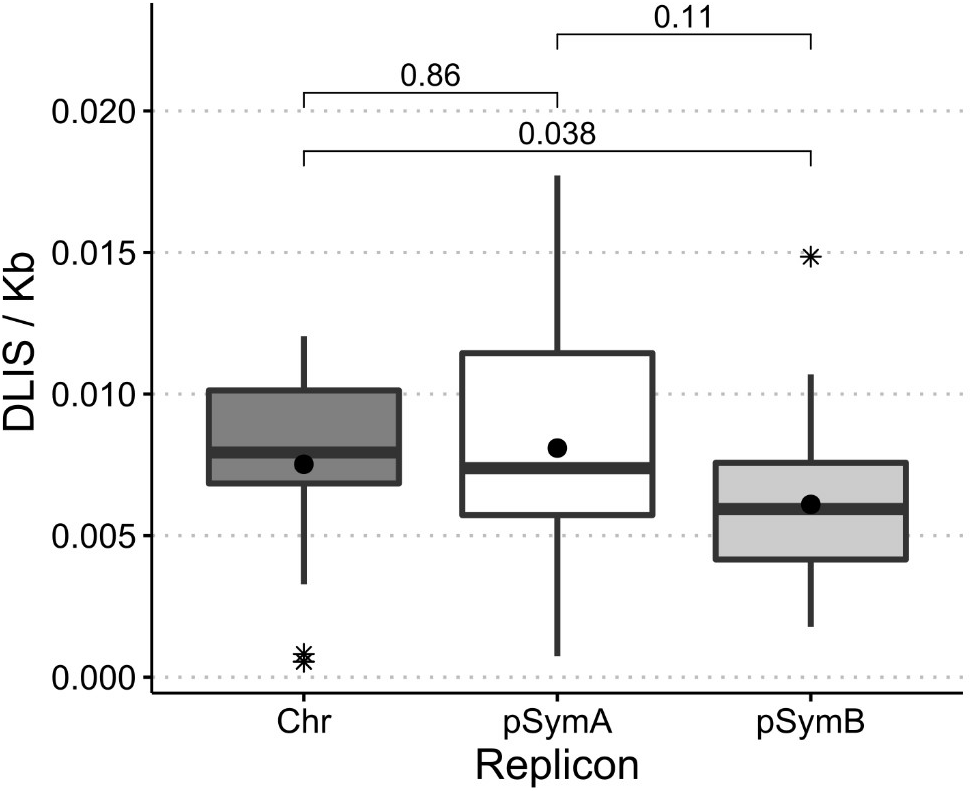
Distribution of DLIS in *E. meliloti* replicons. Box-plot of the number of DLIS per kilobase for the three replicons of *E. meliloti* in the 26 strains used. Chr: Chromosome; pSymA: megaplasmid pSymA; pSymB: megaplasmid pSymB. Asterisks: outliers. Black circles: average. The statistical analysis was performed with R. P-values were calculated using the Kruskal-Wallis test.

In general, the precision was relatively stable (89% on average for DLIS, Supplementary material, Table S4) not showing a significant correlation with the phylogenetic distance. However, a significant correlation of dDDH with the total number of discarded sequences was found (ρ=−0.44,*p=0.032* for all the analysed genomes and ρ=−0.8920158, *p=0.00052* for the complete genomes). In the case of complete genomes, the precision was more variable ranging from 66% to 100%. This could have been due to the presence of regions with consecutive ISs, repeated sequences or rearranged genomic segments, which in complete genomes can generate false positives, while in the case of draft genomes, these regions are present on scaffold ends and usually taken as true DLIS. Another source of false positives was related to mobile group II introns. Mobile group II introns are ribozymes and retroelements present in most *E. meliloti* strains and which natural target site lies within insertion sequences (ISRm2011-2 in the case of RmInt1 and on the left and right inverted repeats of ISRm17 in the case of RmInt2; Toro et al., 2018). To avoid the errors in the detection generated by group II introns, comparisons between *E. meliloti* 1021 and GR4 were run using the -S (Shift) option using values from 100 to 5000. For -S values between 100 and 2000 the results were the same, but with values ranging from 3000 to 5000 the results showed an improvement in the identification of DLIS, including now the partial ISs flanking group II introns (Supplementary material, Fig. S6.B). 30 new DLIS were detected, 23 corresponding to *ltrA* (Group II intron-encoded protein Ltr, a multifunctional protein that promotes group II intron splicing and mobility) flanking ISs and 7 corresponding mostly to multiple consecutive IS. Nevertheless, 4 DLIS that were correctly detected in the normal mode were missing (Supplementary material, Table S6).

In addition, ISCompare was run using as IS database a multifasta file containing all the *ltrA* gene variants found on *E. meliloti* to compare the *E. meliloti* GR4 and 1021 genomes. A surroundingLen of 500 base pairs and a shift of 4000 base pairs were used (Supplementary material, Fig. S6.C). In this comparison, a total of 6 *ltrA* genes were found on *E. meliloti* 1021. 2 *ltrA* genes presented the same location in both strains, with some minor differences. 2 were correctly detected as differentially located, and 2 that were present only on the reference strain were tagged as discarded. This error was produced by differences in the genomic context surrounding *ltrA*.

For *E. meliloti* GR4, 21 *ltrA* genes were detected, 2 were in the same location in both strains, and of the remaining, 6 were discarded because they were located in a different genomic context, 12 were correctly reported by ISCompare as differentially located, and 1 was manually detected in the discarded QIFs. In all cases, the introns that failed to be correctly reported presented differences in the genomic context, especially differentially located ISs.

In addition a comparison of *E. melioti* GR4 with *E. meliloti* G4 (Toro *et al*., 2016) (Supplementary material, Table S1) was done. G4 raw reads were downloaded from SRA (SRR2078187), assembled using SPAdes (Bankevich *et al*., 2012) and annotated with Prokka (Seemann, 2014) at Galaxy Australia web server (Afgan et al., 2016; https://usegalaxy.org.au/). In that case, ISCompare found 6 DLIS, including all the previously reported by Toro et al. (2016), except a difference in a group II intron. This was most likely due to the difficult assembly of regions containing group II introns.

### 3.4 Detection of IS481 within the pertactin gene of *Bordetella pertussis*

*B. pertussis* is the causative agent of whooping cough, which has reemerged as a public health threat despite broad vaccine coverage. The re-emergence of this pathogen has been correlated with the transition from the use of whole-cell pertussis vaccines to acellular component vaccines (Bart *et al*., 2014; Melvin *et al*., 2014) which usually contain up to 5 purified *B. pertussis* antigens, namely the pertussis toxin, (PT), Filamentous Hemagglutinin (FHA), Pertactin (PRN) and Fimbriae (FIM2 and FIM3) (Dewan *et al*., 2020). Among this, the pertactin, a highly immunogenic outer membrane protein that promotes adhesion to tracheal epithelial cells (Inatsuka *et al*., 2010), has been implicated in vaccine-driven evolution presenting different types of knock-out mutations in circulating strains. One of the most common mutations is the insertion of IS481 within its coding sequence (Maruya and Saeki, 2010; Hiramatsu *et al*., 2017; Belcher and Preston, 2015; Zomer *et al*., 2018). Three different locations of IS481 within *prn* have been reported (Pawloski *et al*., 2014). We used the IS481-interrupted *prn* sequences KF804023.1, KC445198.1 and KC445197.1 to look for representative genomes at NCBI Assembly database. Three representative genomes (Table S1.) corresponding to *B. pertussis* strains I127, J412, and J299 were analysed with ISCompare. ISCompare correctly identified an IS481 insertion within the pertactin gene in *B. pertussis* strains J299 and J412, and the IS481 insertion at the 3’-end of strain I127 pertactin gene. In addition, 7 shared DLIS were found in all the genomes (Fig. 4. Black circles and triangles; Table S9). After a manual verification using the graphic report, 31 and 42 DLISs were found for I127 and J299/J412 correspondingly (Fig 4. Grey circles and triangles), mainly involving cases on which two consecutive IS481 were detected in one genome, while only one was found in the other.

**Fig. 4.**
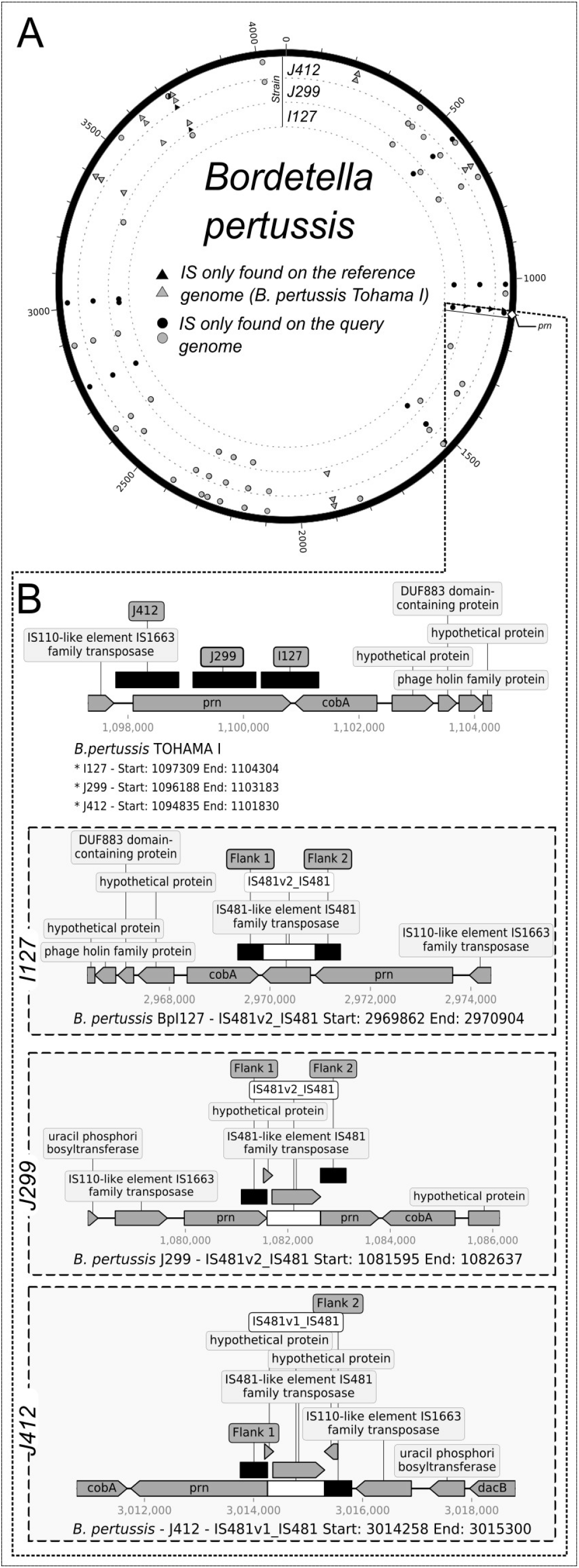
Genomic distribution of DLIS in *Bordetella pertussis* strains with an IS481 insertion within the pertectin gene. A. Circos plot showing the genomic location of DLISs in *B. pertussis. B. pertussis* TOHAMA I was used as reference and compared to strains I127, J299 and J412. B. Adaptation of ISCompare graphic report showing the IS481 insertion within the pertactin gene. Light-grey triangles and circles: DLIS identified by manually analysing the VD category. Black circles and triangles: DLIS category. Circles: DLIS detected in the query genome. Triangles: DLIS detected in the reference genome.

### 3.5 Comparison with ISseeker

Although there are several tools for the comparison of insertion sites between bacterial strains, most of them are based on the soft mapping of short reads from whole genome shotgun sequencing experiments to a reference genome (Breseq, Barrick et al., 2014; Transposon Insertion Finder, Nakagome et al., 2014; ISMapper, Hawkey et al., 2015; panISa, Treepong et al., 2018; and ISSeeker, Adams et al., 2016). The only one with features similar to those of ISCompare is ISSeeker.

To compare the performance of ISCompare with ISseeker we analysed the location of the ISRm2011-2 on *E. meliloti* GR4 and 1021 using both programs (Fig. 5.; Supplementary material, Table S7). ISseeker outputs a table with the mapping coordinates of left and right query IS flanks relative to the reference genome. In addition, it attempts to find mates by comparing the mapping coordinates of all the flanks. Thus, for the analysis of ISSeeker results, we considered as DLIS those ISs which flanks were correctly mapped and mated. All the reported flanks and ISs were manually analysed to define whether they were DLIS or not. Our program performed similar to Isseeker (Fig. 5A), both correctly detecting 8 DLIS, and several ISs located in a different genomic context. However, ISSeeker reported ISs which were discarded by ISCompare because they contained flanks that produced blastn matches with multiple genomic regions, and in most cases corresponded to false positives. In addition, ISSeeker failed to detect *ltrA* related DLIS (15 ISs) which were detected by ISCompare using the shift option (Fig. 5A, Gray bars).

**Fig. 5.**
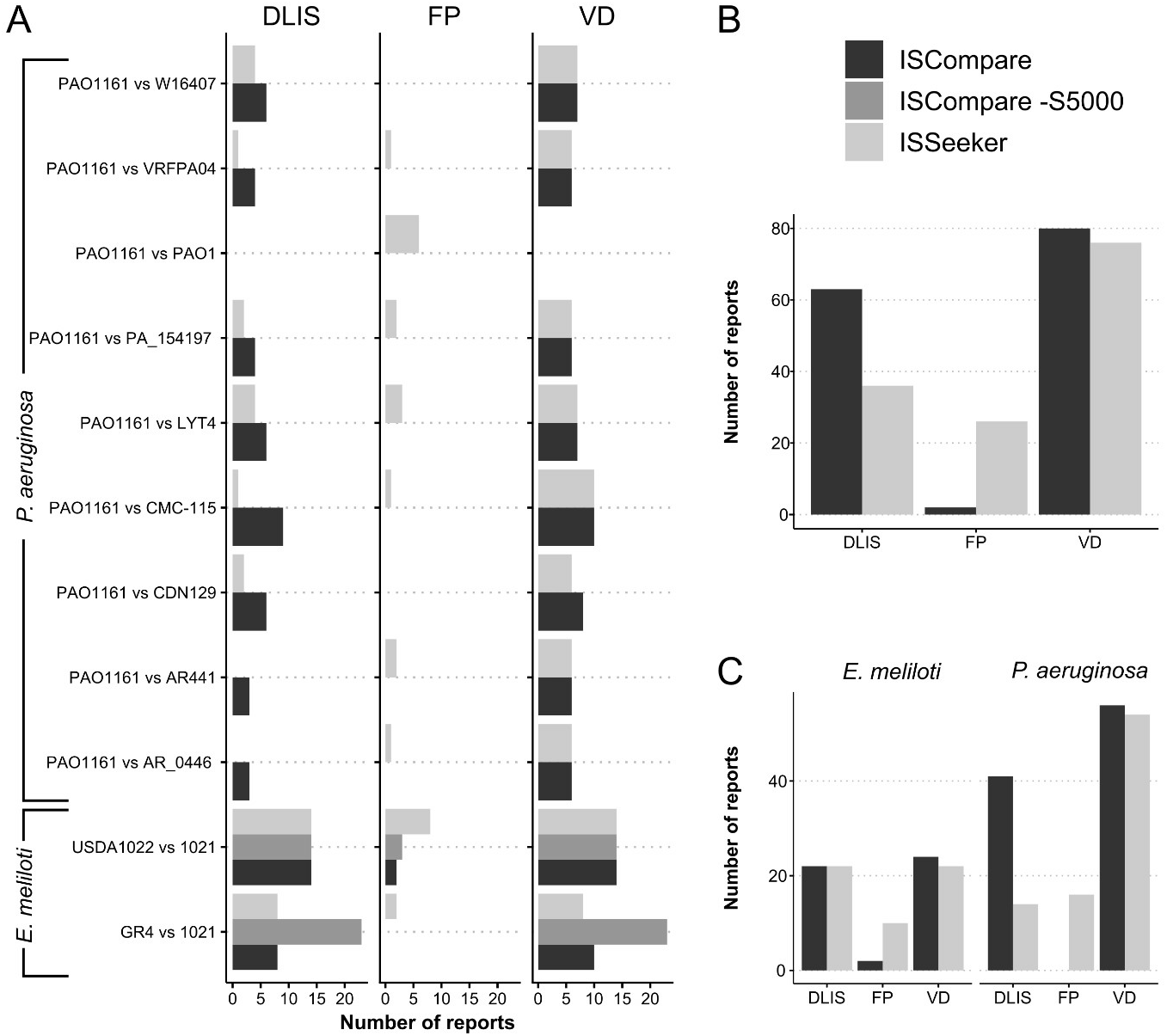
Performance of ISCompare in comparison with ISSeeker. ISCompare was compared with ISSeeker using selected ISs from *E. Meliloti* and *P. Aeruginosa* strains. A. Number of DLIS, VD and FP reports for *P. Aeruginosa* and *E. Meliloti* strains. B. Total number of DLIS, VD and FP reports. C. Total number of DLIS, VD and FP reports discriminated by specie. VD: Total identified DLIS, including those that were identified manually. FP: false positives. DLIS: Differentially located ISs reported. In the case of ISSeeker, only ISs with assigned mates were considered as DLIS. Black: ISCompare; Grey: ISCompare with -S 5000 option; Light-gray: IS-Seeker.

We also tested both programs for the identification of DLIS in the case of draft genomes. We tried to run ISSeeker for the comparison of *E. meliloti* USDA1022 (157 scaffolds) with *E. meliloti* 1021. In that case, an error occurred and ISSeeker code had to be modified (ContigBlastHit.pm; Supplementary material, file S1.) to include ISs found on scaffold ends. The results were fairly comparable for both programs (14 DLIS found), with ISCompare producing fewer false positives (Fig. 5A.; Supplementary material, Table S7). We also tested ISSeeker with the *P. aeruginosa* dataset. In that case, the performance of ISCompare was better than that of ISSeeker, and less false positives were produced (Fig. 5A and C.; Supplementary material, Table S8). From these results we can conclude that ISCompare performed better than ISSeeker (Fig. 5B.), achieving a slightly better sensitivity and precision.

## 4 Discussion

ISCompare is a tool for identification of DLIS which uses blast (Altschul *et al*., 1990) for the pairwise genomic comparison of ISs location. ISCompare requires two genomes to compare in an annotated genbank format and an optional IS database. The output consists of several files, being the final results table and the graphic report the most valuable ones. The final results table, reports all the differentially located IS candidates classified in 8 categories (Supplementary material, Fig. S5). First is the ‘DLIS’ category which has the highest confidence, then categories where there were consecutive IS or genes with multiple copies which can produce false positives, and finally categories where the IS were discarded from the analysis because of different reasons (i.e. flanks not found on the reference genome, flanks with query coverage below the cutoff values, etc). The graphic report file contains the genomic context of both the reference and query genomes, providing the means for a rapid inspection and verification of the results.

We analysed the sensitivity and precision of ISCompare using artificially generated genomes with randomly generated insertions of a specified IS, showing that ISCompare performed very well, achieving high precision 100% (92) and sensitivity 94% (99.1) for DLIS (VD) categories.

When we performed the analysis with real genomes, an increment in the number of false positive cases was observed (precision of 89% for *E. meliloti* strains). In that case, many of the false positives reports corresponded to DNA regions where the context of one side of the IS is correct, but the other is different, in general, because of genome rearrangements. Other false positives corresponded to regions containing very similar paralogous genes, integrases, which might be phage related, and in the case of *E. meliloti*, to group II introns. Interestingly, some of the locations reported presented a change of one IS for another. It should be noted that some IS might be absent on ISFinder database (i.e. ISRm19), in such cases the ISs present on the strains to analyse should be previously detected and added to IS.fasta file.

A feature of ISCompare is that it can be also used to analyse the genomic context of transposon, phages or any other gene or DNA sequence in general. As a demonstration, on this work we used ISCompare to study the differentially located Group II introns in *E. meliloti*. For this comparison the -S shift option, an exclusive running mode of ISCompare, greatly improved the detection of DLIS. From all the analysed cases, our recommendation is to run ISCompare in normal mode, and if regions with many consecutive IS or repeated sequences (such as group II introns) are present, run also in the shift (-S) mode. In both cases the DLIS results are very confident and when combined should improve the sensitivity of the program.

Finally we want to address the differences between ISCompare and ISSeeker. Both programs are based on the search of IS flanks from a query genome on a reference genome, however, there are several differences. First, ISSeeker presents the output in a different way, since it was thought for the fast and high-throughput comparison of the genome of many strains to a single reference genome. On the other hand, ISCompare was designed for pairwise IS location comparison. Second, the output table of ISSeeker shows the flanks of the detected IS on the query genome, with the nearest position where those flanks align on the reference genome, and the position of all the IS found on the reference genome. The identification of the DLIS can then be manually or graphically done by comparing those values. ISCompare, instead, generates a table containing the identified DLIS not requiring further analysis. Third, it should be noted that to get a complete idea of all the relocated ISs, ISSeeker should be run a second time using the reference genome as query. This is required since for divergent enough strains, ISs could be present on genomic regions exclusively found in the reference strain. ISCompare focuses on identifying DLIS on conserved genomic regions, in the hope to find DLIS responsible for phenotype changes. Fourth, ISSeeker filtering steps are based on percent identity cut-offs, while ISComare uses query coverages and E-value cut-offs. Fifth, ISseeker requires previous knowledge of an IS present in both genomes while ISCompare can be used to look for DLIS using a complete (or partial) database of ISs (such as ISFinder). Further, using -I option, ISCompare will use ISFinder (Siguier *et al*., 2006) blast server to look for and download all the IS families found on the query and reference genomes. Last, although ISSeeker is faster, ISCompare runs fast enough and outputs several tables with information about the ISs found on the query and reference genomes, cases of consecutive IS which might yield false positives, a clear annotation of CDS flanking the insertion sites, and a graphical PDF report showing the IS surroundings for manual verification of DLIS status.

For all this, we think that ISCompare is a program that provides an easy and straightforward approach to look for DLIS between a pair of related bacterial genomes.

## Acknowledgements

This work was supported by the National Science and Technology Research Council (Consejo Nacional de Investigaciones Científicas y Técnicas—CONICET, Argentina) (PIP 2014-0420), the Ministry of Science Technology and Productive Innovation (Ministerio de Ciencia Tecnología e Innovación Productiva—MinCyT, Argentina) (PICT-2016-0171, PICT-2015-2452) and the National University of La Plata (Universidad Nacional de La Plata). M. J. L. is supported by the CONICET, N. A. and E. G. M. are supported by grants from the MinCyT.

## Author contributions

M. J. L. wrote the ISCompare program, interpreted data, and edited the figures and manuscript; E. G. M. ran simulations, evaluated the performance of ISCompare and interpreted data; N. A. evaluated the performance of ISCompare and interpreted data. All authors approved the final version of the manuscript.

## Supplementary material

**Fig. S1.**
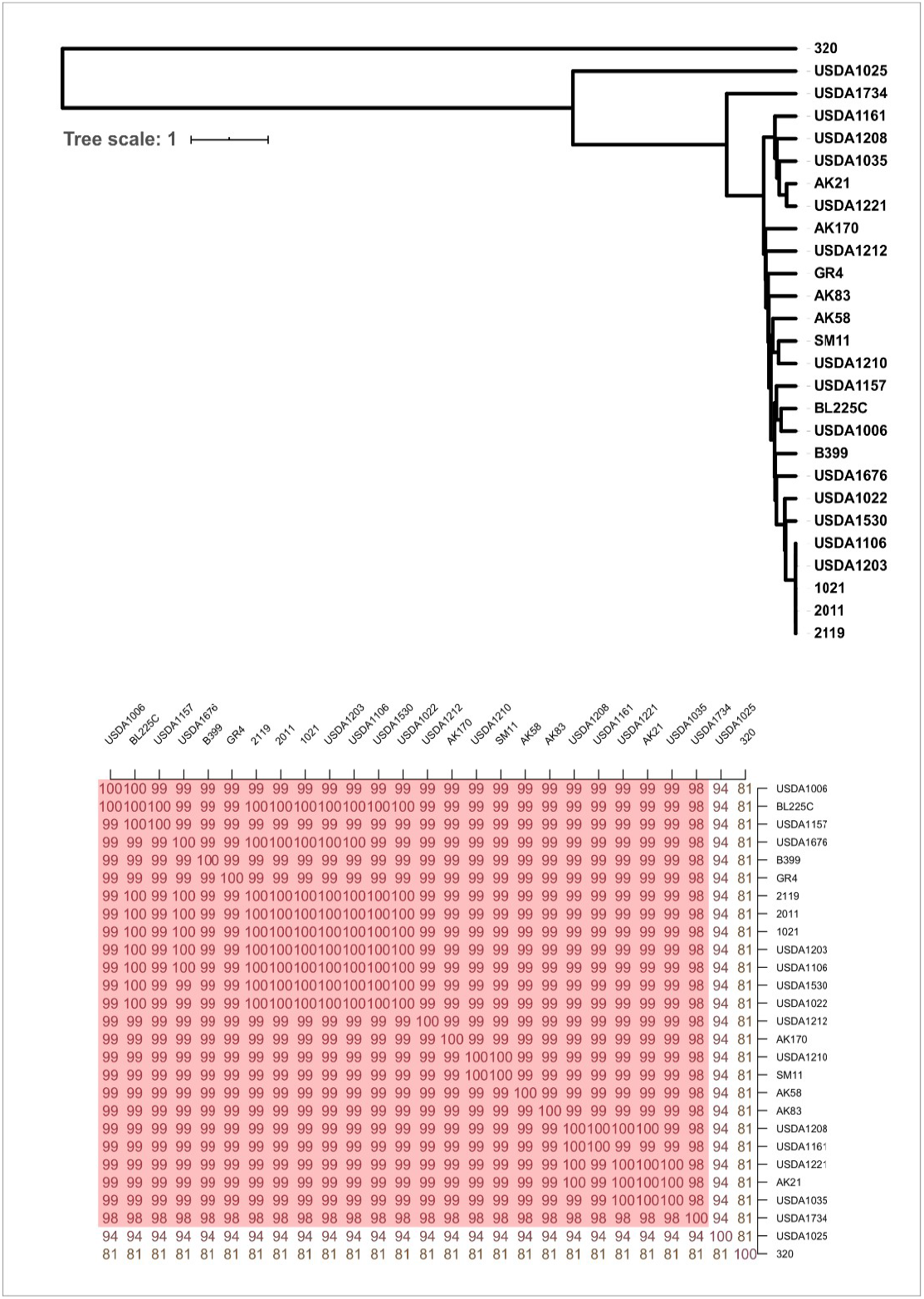
*E. meliloti* Average Nucleotide Identity (ANIb) matrix and UPGMA distance tree. The ANIb matrix and UPGMA distance tree were calculated using the ANI-matrix calculator at Kostas lab server (http://enveomics.ce.gatech.edu/g-matrix/). The accession numbers of the *E. meliloti* genomes used are listed on Table S1.

**Fig. S2.**
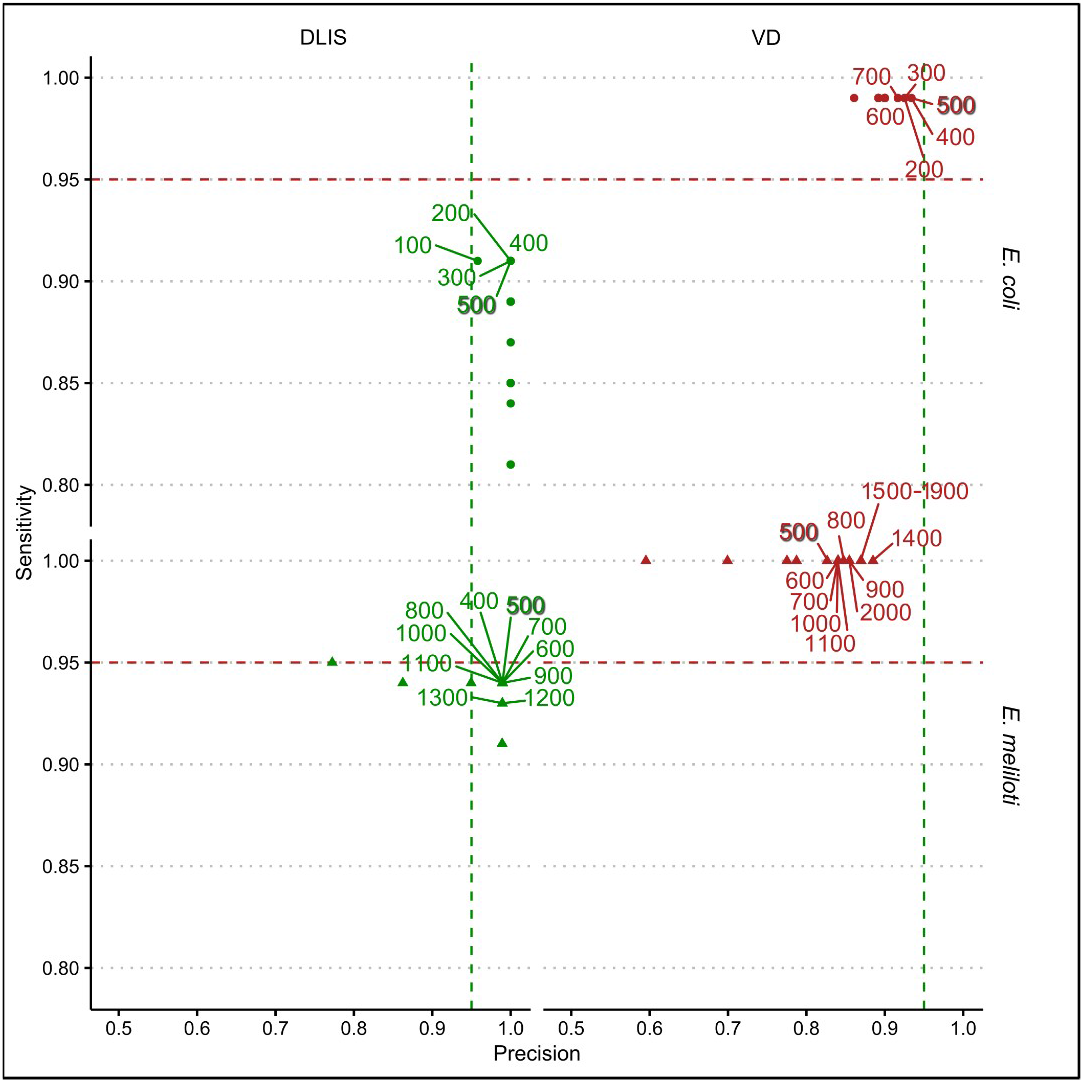
Sensitivity and precision of ISCompare using different SurroundingLen values. SurroundingLen parameter optimization was evaluated using a range of nucleotide lengths between 100 and 2000. *E. coli* K-12 substr. MG1655 genome was compared with an artificial genome of the same strain containing 100 IS30 random insertions. In the case of *E. meliloti*, the comparison was done between strain 1021 as reference genome and an artificial *E. meliloti* 2011 genome with 100 ISRm5 random insertions as query. DLIS: Differentially located ISs; VD, discarded cases or cases tagged for manual verification.

**Fig. S3.**
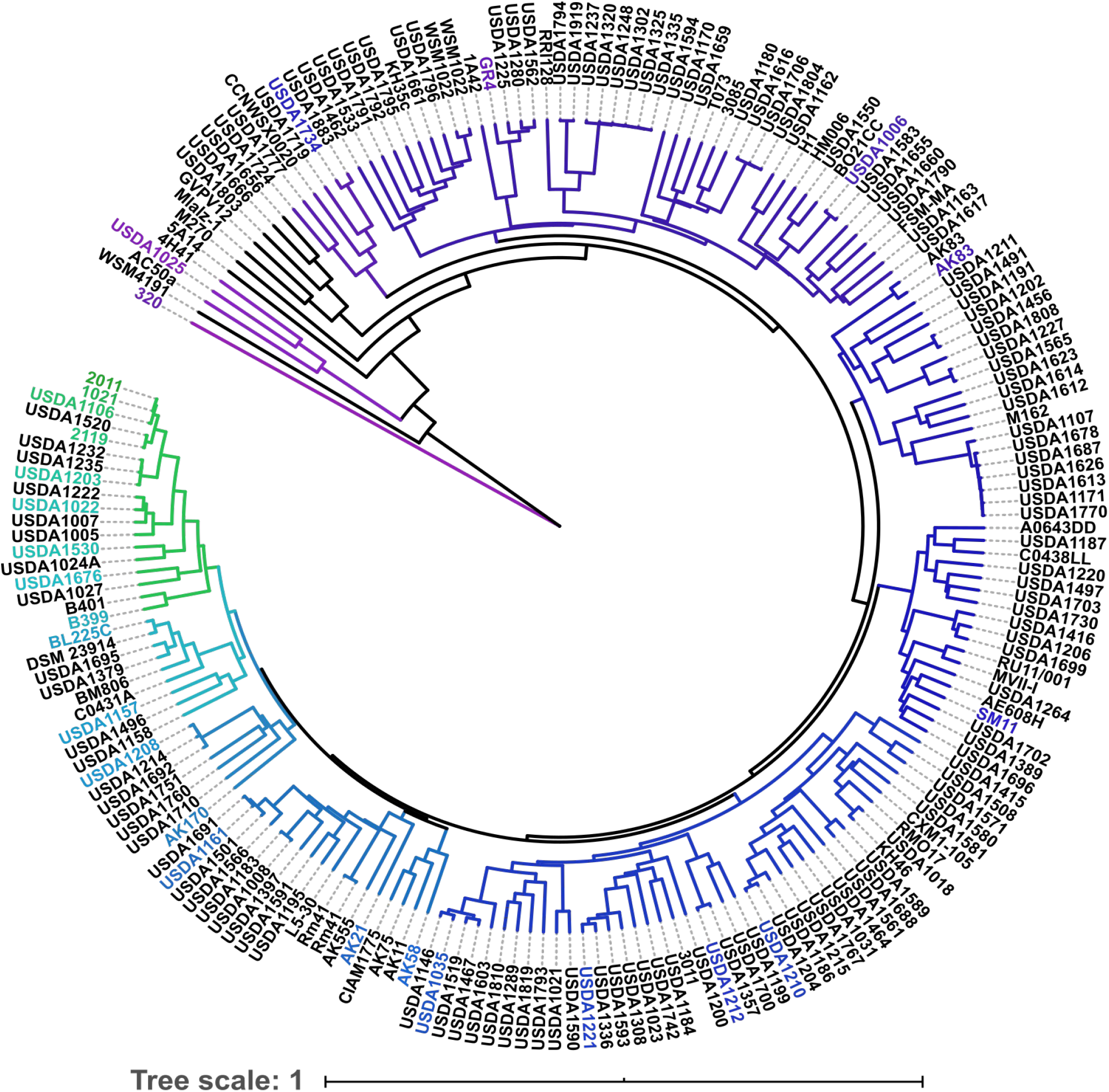
Phylogenetic tree of all the sequenced *E. meliloti* strains at NCBI genomes database. The phylogenetic tree was downloaded from NCBI genomes database. *E. meliloti* strains were selected according to their phylogenetic distance to the reference strain 1021. Selected strains are shown in color, from green for nearly related strains, to purple for more distant strains. The phylogenetic tree was edited using ITOL server (Letunic and Bork, 2019, https://itol.embl.de/).

**Fig. S4.**
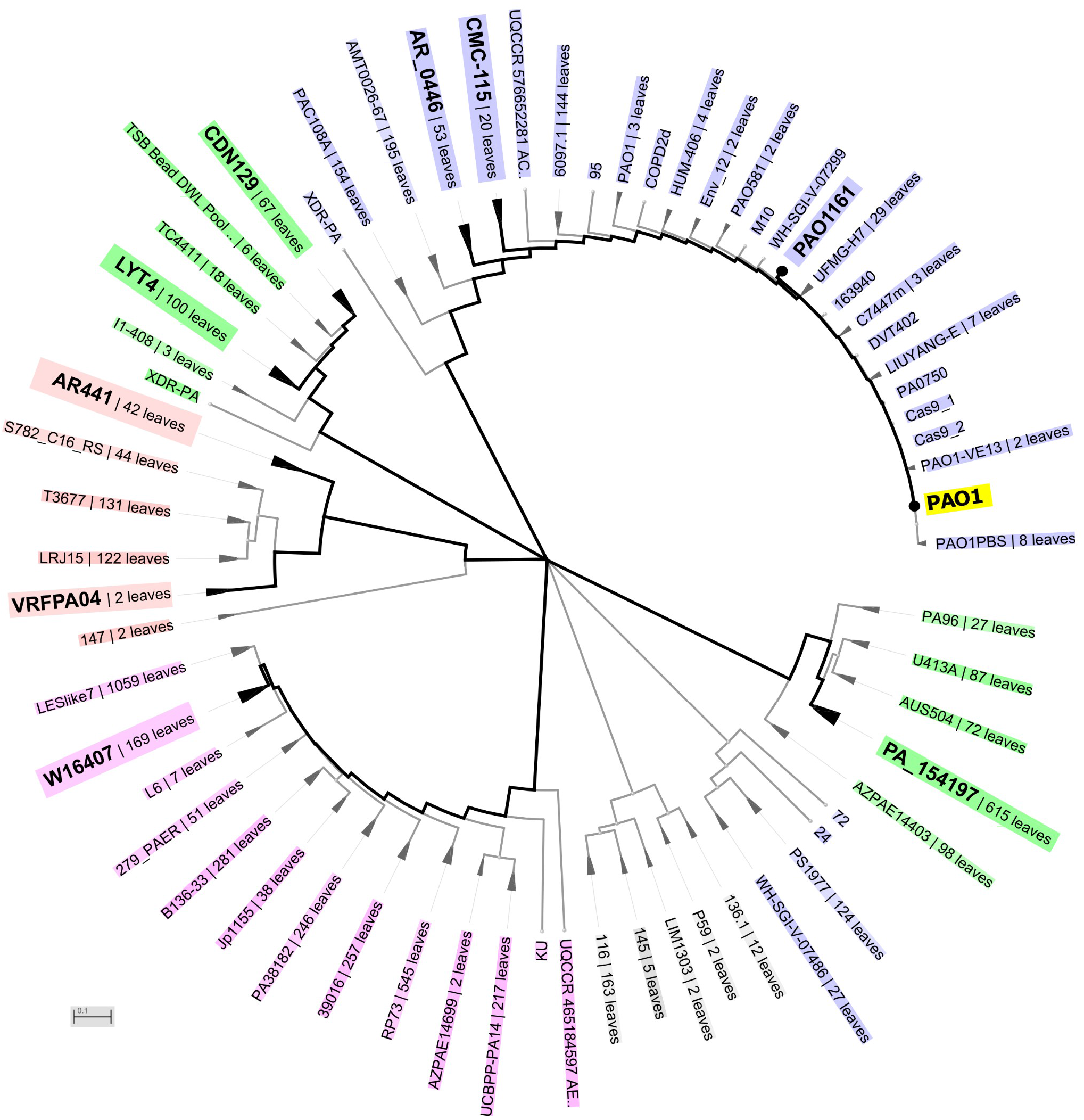
Phylogenetic tree of all the sequenced *P. aeruginosa* strains at NCBI genomes database. The phylogenetic tree was downloaded from NCBI genomes database. *P. aeruginosa* strains were selected according to their phylogenetic distance to the reference strain PAO1. Selected strains are shown in bold fonts. Collapsed branches are displayed as triangles. Leaves are shown as dots. The phylogenetic tree image was manually edited with Inkscape.

**Fig. S5.**
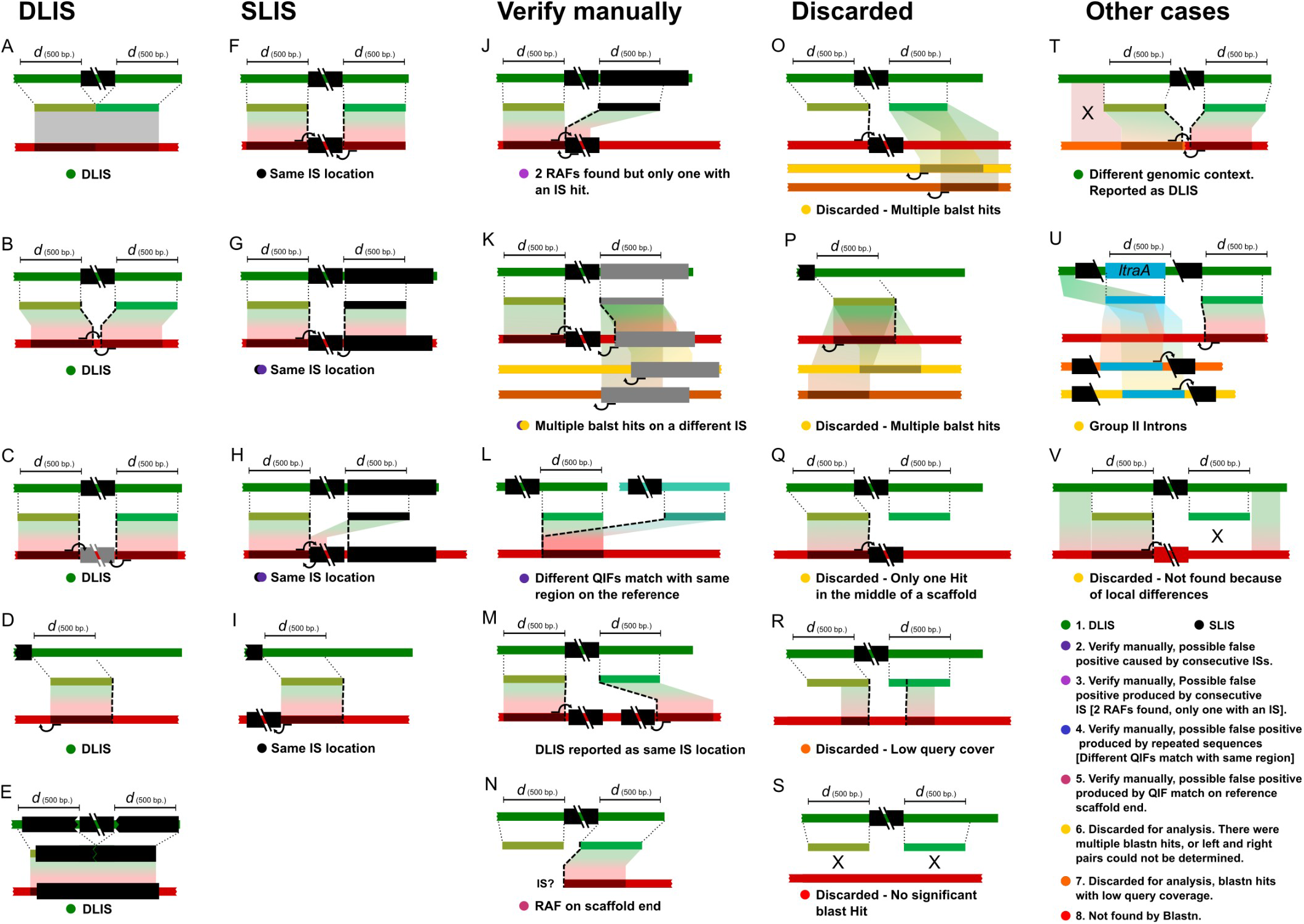
Schematic representation of the possible cases resulting in DLIS, SLIS, and VD reports. Panes A-E represent cases which would be correctly identified by ISCompare as DLIS. Panes F-I correspond to cases which will be reported only using the -rs (report SLIS) option. Panes J-N are cases reported for manual verification, most of them involving repeated sequences. Some of these cases could be DLIS. Panes O-S are cases discarded from the analysis due to non significant or multiple blastn hits. Panes T-V, are other particular cases which could produce false positives. The query and reference genomes are represented by thick lines (green for query, and red/orange/yellow for reference) and ISs are represented as black boxes. Grey boxes represent different ISs and other genes. *ltrA*: group II intron-encoded protein LtrA. Colored circles indicate the category of differentially located IS candidate (Green, DLIS; Black, SLIS; Purple; magenta and dark blue are “Verify manually” categories; Yellow, orange and red are “Discarded from the analysis” categories).

**Fig. S6.**
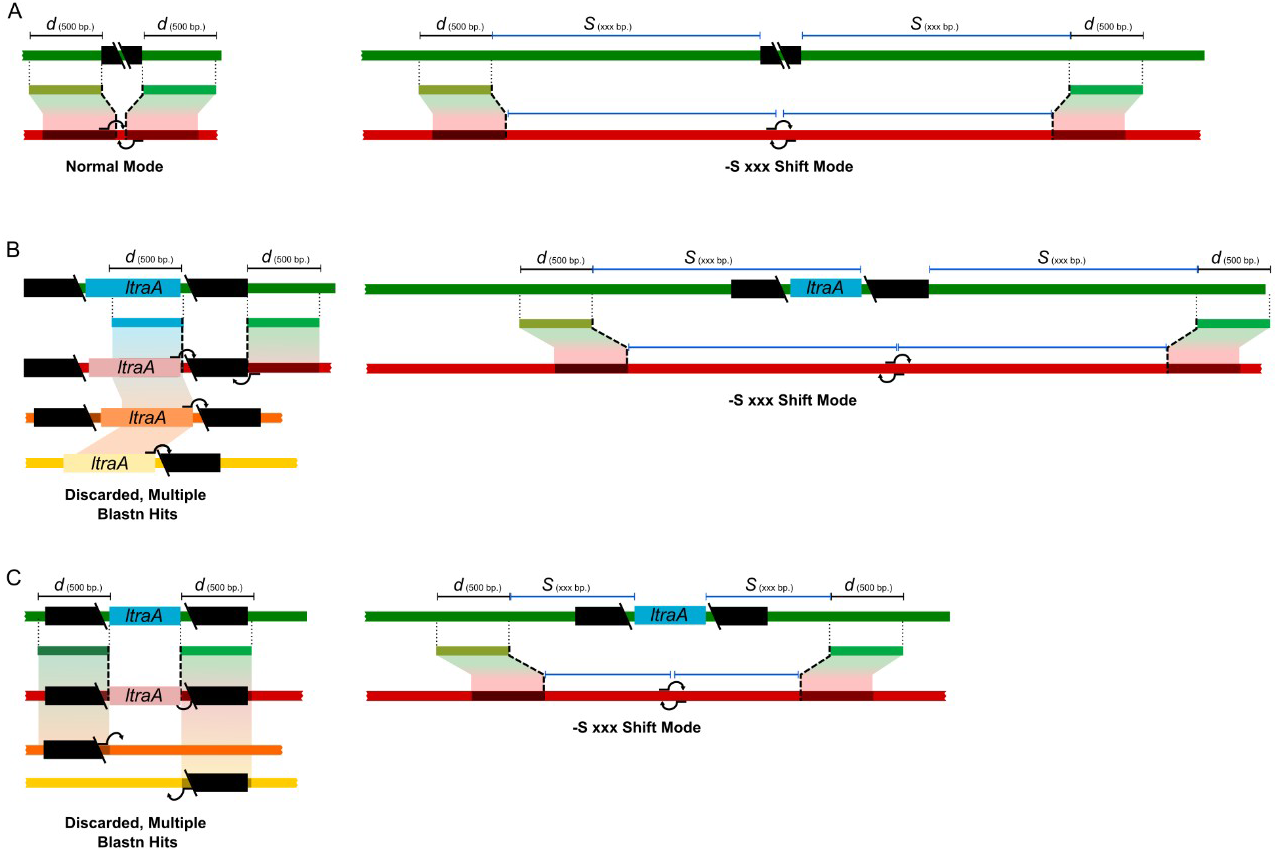
ISCompare shift mode. A. Schematic representation of the algorithm variations in the -S shit mode. B. Example of usage of -S mode for the identification of differentially located ISs flanking Group II introns. C. Example of usage of -S mode for the identification of differentially located Group II introns using *ltrA* gene.

## Supplementary Tables

**Table S1. Genomes used for ISCompare evaluation**

**Table S2. SurroundingLen parameter optimization.** Sheet 1, results of the comparison of *E. coli* K-12 substr. MG1655 with an artificial genome of the same strain containing 100 IS30 random insertions. Sheet 2, results of the comparison using *E. meliloti* strain 1021 as reference genome and an artificial *E. meliloti* 2011 genome with 100 ISRm5 random insertions as query. Sheet 3, Sensitivity and precision analysis. TP, true positives, FP, false positives, TP*, true positives manually found by inspecting the VD reports.

**Table S3. ISCompare evaluation using 3,000 random IS30 insertions.** Sheet 1: Statistical analysis. Sheet 2: Location of 3,000 randomly inserted IS30. Set 1. Sheet 3: ISCompare results using a compilation of IS. Sheet 4: Location of 3,000 randomly inserted IS30. Set 2. Sheet 5: ISCompare results using IS30 as query IS. TP, true positives, FP, false positives, TP*, true positives manually found by inspecting the VD reports.

**Table S4. Analysis of differentially located ISs on *P. aeruginosa* strains.** Sheet 1: ISCompare result analysis. TP, true positives, FP, false positives, TP*, true positives manually found by inspecting the VD reports. Sheet 2: ANIb and DDH results.

**Table S5. Analysis of differentially located ISs on *E. meliloti* strains.** Sheet 1: ISCompare result analysis. TP, true positives, FP, false positives, TP*, true positives manually found by inspecting the VD reports. Sheet 2: ANIb calculated using ANI matrix calculator server. Sheet 3: dDDH results from http://ggdc.dsmz.de/ggdc.php.

**Table S6. Comparison of ISCompare results using the normal vs the Shift mode (-S).** A comparison of DLIS in *E. meliloti* 1021 and GR4 strains was done using ISCompare with -S option set to 5,000 nucleotides.

**Table S7. Comparison of ISCompare and ISSeeker using *E. meliloti* genomes.** *E. meliloti* 1021 was used as reference, and compared to GR4 and U1022 strains as queries. In the case of GR4 strain, the -S option was also evaluated. As ISSeeker only can analyse one IS at a time, only ISRm2011-2 location was analysed using both programs. Sheet 1: ISCompare vs ISSeeker results summary. Sheet 2: ISSeeker, 1021 vs GR4 results. Sheet 3: ISCompare, 1021 vs GR4 results. Sheet 4: ISCompare with -S 5000 setting. 1021 vs GR4 results. Sheet 5: ISSeeker, 1021 vs USDA1022 results. Sheet 6: ISCompare. 1021 vs USDA1022 results. Sheet 7: ISCompare with -S 5000 setting. 1021 vs USDA1022 results.

**Table S8. Comparison of ISCompare and ISSeeker using *P. aeruginosa* genomes.** Sheet 1: ISCompare vs ISSeeker results summary. Sheet 2: ISSeeker results for all the analysed *P. aeruginosa* strains. Sheet 3: results for all the analysed *P. aeruginosa* strains.

**Table S9. Comparison of *B. pertussis* TOHAMA I with strains I127, J299 and J412 containing a IS481 insertion on the pertactin autotransporter gene.** Sheet 1: Summary. Sheet 2: Results of TOHAMA I vs I127 using ISCompare. Sheet 3: Results of TOHAMA I vs J299 using ISCompare. Sheet 4: Results of TOHAMA I vs J412 using ISCompare.

